# LINbase: A Web service for genome-based identification of microbes as members of crowdsourced taxa

**DOI:** 10.1101/752212

**Authors:** Long Tian, Chengjie Huang, Lenwood S. Heath, Boris A. Vinatzer

## Abstract

The development of next generation and third generation DNA sequencing technologies in combination with new efficient algorithms allows scientists to economically, quickly, and precisely identify microbes at all taxonomic levels and even attribute pathogen isolates to specific disease outbreaks. However, current taxonomic practice has not kept up with the sequencing revolution and continues to rely on cumbersome journal publications to describe new species. Here we introduce a Web service that allows any user to genomically circumscribe any monophyletic group of bacteria as a taxon and associate with each taxon a name and short description. Any other user can immediately identify their unknown microbe as a member of any of these crowdsourced taxa using gene or genome sequences. The Web service is called LINbase. It leverages the previously described concept of Life Identification Numbers (LINs), which are codes assigned to individual organisms based on genome similarity. Most genomes currently in LINbase were imported from GenBank, but users have the option to upload their own genome sequences as well. Importantly, LINbase allows users to share the precise identity of their sequenced genomes without sharing the actual genome sequences, making not yet published or private genome sequences discoverable by the scientific community stimulating collaboration between academia and industry. LINbase is available at http://www.LINbase.org.

## Introduction

Fast and precise pathogen identification is crucial in human, animal, and plant disease diagnosis to identify the most effective treatment and to limit disease spread [1]. Precise identification of microbes is also important in many other fields, for example, when regulating commercial probiotics for human consumption [2] or biopesticides to control plant diseases in agriculture [3]. While we often associate the process of identification with giving an unknown organism a name, the ultimate goal of identification is to predict the characteristics of the unknown organism independently of what its name is, for example, to answer a question such as: does the unknown microorganism cause a certain disease in a specific animal species? The prerequisite for such precise identification is precise classification [4]. Only if microbes are classified into groups (called taxa) in which all members are derived from a most recent common ancestor (MRCA) (*i*.*e*., constitute a monophyletic group) and share a phenotype absent from organisms outside of that same taxon can identification of an unknown as a member of such a taxon lead to precise prediction of its phenotype.

Before the advent of DNA sequencing, classification and identification necessarily relied on phenotypic tests. Therefore, taxa were restricted to groups of microbes that could be phenotypically distinguished from other microbes based on relatively simple lab-based assays [5]. Classification and identification then transitioned to more precise gene-based methods, in particular, sequencing of the 16S rRNA gene [6]. With the development of ever faster and cheaper high throughput DNA sequencing methods, the entire genome of organisms can now be used to classify organisms into monophyletic groups and identify them as members of these groups based on identification of single nucleotide polymorphisms (SNPs) [7], construction of phylogenetic trees based on conserved genes [8, 9], or measures of genome similarity at the whole genome level expressed as average nucleotide identity (ANI) [10]. Conceptually, a taxon can now consist of microbes that share nothing other than a single mutation inherited from their most recent common ancestor compared to organisms outside of that taxon. If that single mutation changed the phenotype of the microbes belonging to the taxon, then identifying an unknown as a member of that taxon could predict the phenotype of the unknown. Or, in another example, if all microbes with a specific SNP were isolated during a specific disease outbreak, identifying an unknown as a member of the corresponding taxon would be informative of a transmission event and would become epidemiologically important to stop the further spread of a disease.

However, the fundamental unit of current taxonomy is not the smallest distinguishable unit based on genome sequencing, but it is the species, whereby each named species is associated with a “type” strain, which is considered the name-bearing strain of the species [11]. Since current taxonomy is grounded in microbiological history, the valid publication of a new named microbial species requires much more than sequencing the genome of a type strain and reporting a distinctive phenotype. Besides showing that the type strain of the newly named species has less than 95% ANI compared to genomes of type strains of already named species, valid publication requires a long list of results derived from laboratory-based phenotypic tests [12]. Moreover, the process of validly publishing a named species still relies on publication of a traditional manuscript even though the key genomic and phenotypic information of a new species could be easily reduced to a simple database entry, similar to what has been proposed for the Digital Protologue Database [13]. Another limitation with using the species as the smallest unit of bacterial taxonomy is that members of the same species sometimes still vary considerably in regard to some phenotypes, for example, a single plant pathogen species may include many different strains with many of them having a different host range [14]. Although classification schemes at intraspecific levels exist that take into account phenotypic differences between strains belonging to the same species, they are not consistent across species, making it difficult to interpret identification results based on a particular scheme for scientists who do not have familiarity with a particular species-specific scheme.

To address the above-listed limitations of current taxonomy and to take full advantage of genome sequencing for precise classification, the Life Identification Number (LIN) system was introduced [15, 16]. The LIN system classifies bacteria based on reciprocal ANI. In its current implementation, LINs consist of 20 positions, each representing a different ANI threshold. ANI thresholds range from 70% at the left-most position to 99.999% at the right-most position (**Figure 1**). Importantly, LINs are assigned to individual genomes, whereby genomic relatedness between genomes is represented by the length of the longest common prefix of their LINs: the longer the LIN prefix is that is shared by two genomes, the more similar the genomes are to each other. To assign a LIN to a newly sequenced genome, the most similar genome that already has a LIN is identified in a database of genomes and the LIN of the new genome is computed based on its ANI to that most similar genome [17].

**Figure 1.**
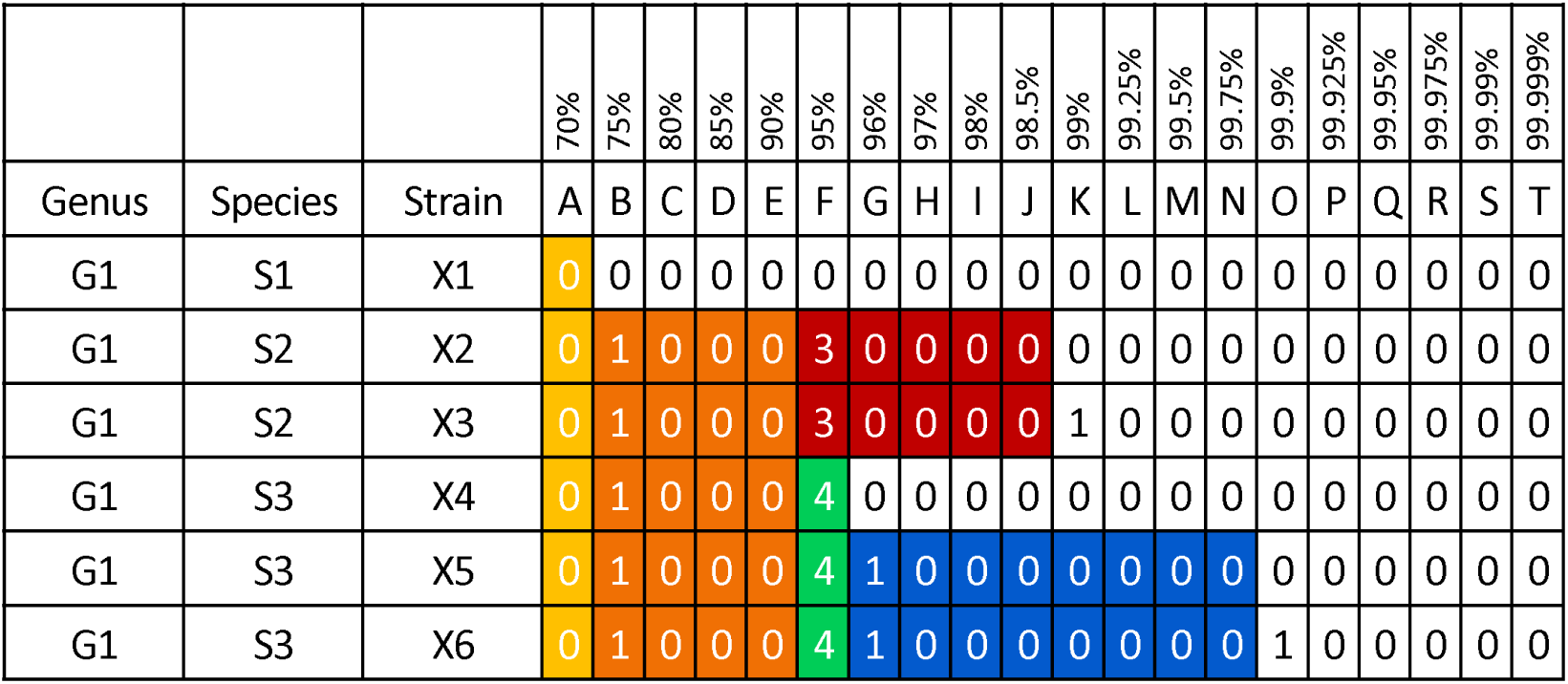
The LIN and LINgroups concept. Each LIN position (A - T) represents an ANI threshold ranging from 70% at position A to 99.999% at position T. LINgroups are used to describe groups of microbes sharing the same phenotype(s). LINgroups are denoted by their shared LIN prefix (from position A to the right-most position that all members of a LINgroup share). For example, using the ANI speciation threshold of 95%, LINgroup 0_A_1_B_0_C_0_D_0_E_3_F_ corresponds to the species G1 S2, and LINgroup 0_A_1_B_0_C_0_D_0_E_4_F_ corresponds to the species G1 S3.

Any group of bacteria that share a LIN prefix of any length is called a LINgroup [18]. If the members of a LINgroup share a phenotype of interest, that single phenotype can be associated with that LINgroup. Therefore, if a microbe is identified as a member of a LINgroup based on its genome sequence, the unknown can be inferred, with high likelihood, to have the same phenotype as all the other members of that LINgroup. Validly published named species and genera can also be correlated with LINgroups. For example, if all known members of a named species share a certain LIN prefix, for example, 0_A_1_B_0_C_0_D_0_E_4_F_, then the LINgroup 0_A_1_B_0_C_0_D_0_E_4_F_ can be associated with that named species and unknowns can be precisely identified as a member of that named species based on their genome sequence (**Figure 1**).

Here we introduce LINbase, a Web service that implements the LIN and LINgroup concepts using an SQL database, efficient algorithms, and an intuitive Web site. Registered users can genomically circumscribe LINgroups and associate them with any phenotype based on their subject knowledge. Users can also associate a LINgroup with any validly published named species or genus based on their taxonomic expertise. This crowdsourcing approach is expected to provide precise genome-based circumscriptions and phenotypic descriptions of taxa, *i*.*e*., LINgroups. To precisely identify microbes, users can query LINbase with genome sequences to determine if an unknown microbe is a member of any circumscribed LINgroup. Users can also upload their own genome sequences to LINbase. Importantly, genome sequences are not shared with other users, but the assigned LINs reveal their precise similarity to all other genomes in LINbase and make them discoverable by all other users, allowing even industry to share their repertoire of genomes without having to share actual DNA sequences. LINbase is fully functional but improvements in regard to speed, resolution, and functionality are ongoing.

## Web Service infrastructure

## Web service

LINbase is built with the LAMP (**L**inux, **A**pache server, **M**ySQL and **P**HP) stack with a RESTful API written in JavaScript and a job scheduler written in the Go programming language. All code is structured in an MVC (Model-View-Controller) framework named CodeIgniter. The analytical parts of LINbase are written in Python. The server currently runs on an Intel Xeon 16-core CPU at 1.90GHz, with 64GB RAM and the CentOS 7 operating system. The Web site can be accessed at http://linbase.org.

## Database management

MySQL 5.6 is used to manage the database and store all relevant metadata. The schema is shown in **Figure 2**. Each table has a primary key and is connected to other tables with a foreign key. There are 4 main tables storing data related to uploaded genomes: the genome table stores the locations of the genome assemblies on the server, the taxonomy table stores the taxonomic information, the MetadataValue table stores associated metadata, and LINs of all uploaded genomes are recorded in the LIN table. The remaining tables serve the purpose of smoothing the data transfer and task management of LINbase. All tables are indexed for optimized query speed.

## Summary of LINbase Functions

LINbase assigns LINs to microbial genomes uploaded by LINbase administrators and users. Users are encouraged to describe microbial taxa that correspond to genera or species or intraspecific groups with distinctive phenotypes as LINgroups. Users can also comment on LINgroups described by other users, search genomes and LINgroups by keyword, and query LINbase using gene sequences and genome sequences.

**Figure 2.**
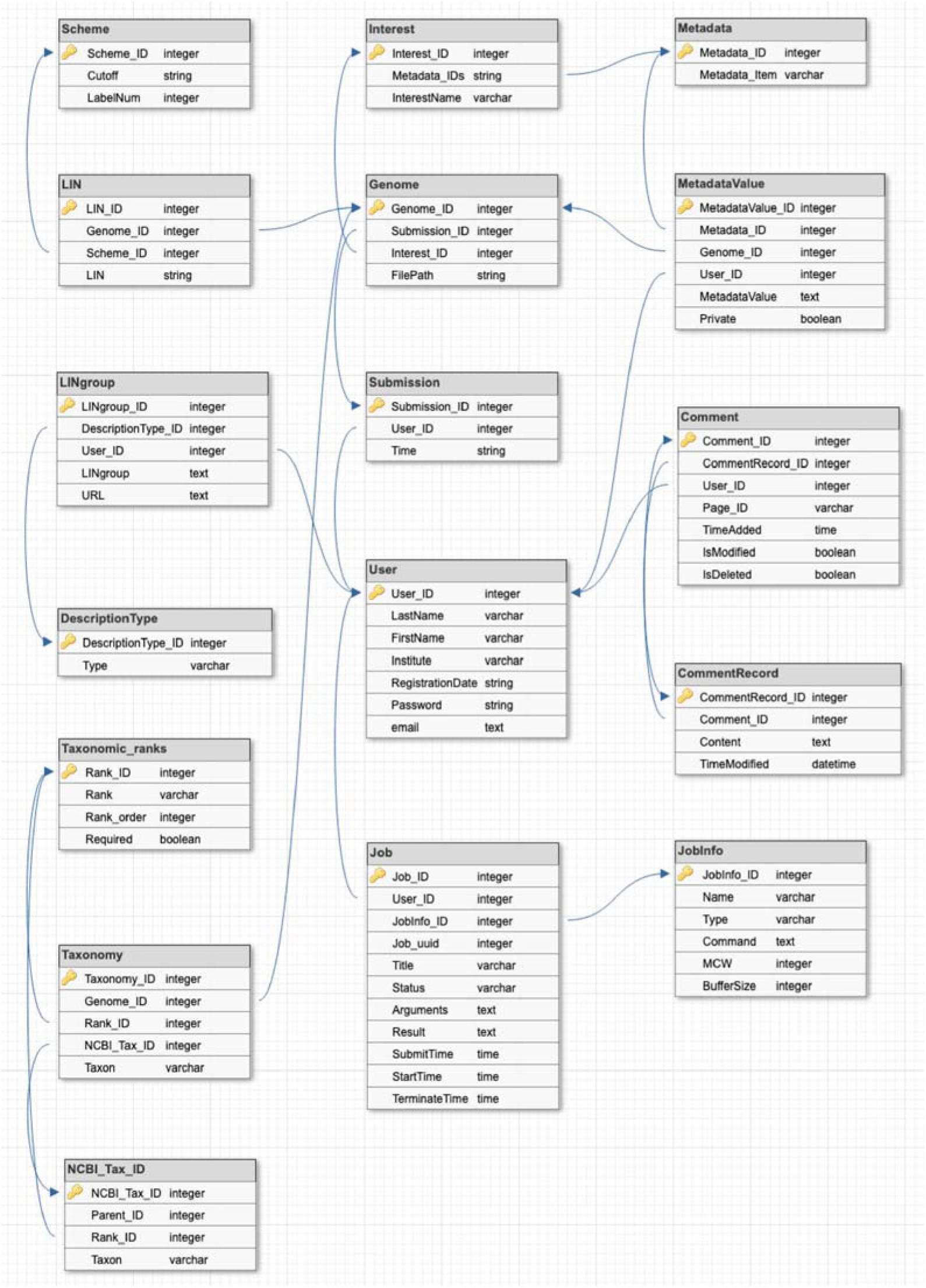
Database schema of the SQL database of LINbase. All relevant data of LINbase are saved in a relational database with the shown schema. Each table is connected with other tables with a primary key and foreign key(s). The first row of each table is the primary key, and the arrow pointing to another table indicates the connection with the foreign key.

At the time of writing, 6204 bacterial genomes have been uploaded to LINbase and 4079 of them have been identified as members of 50 circumscribed LINgroups.

### Genome upload function

The goal of LINbase administrators is to add all microbial genome sequence assemblies of NCBI’s Genbank database to LINbase as long as assemblies satisfy minimal quality standards, such as having fewer than 500 contigs. However, if users are interested in Genbank genome sequences that have not been added to LINbase yet, they can upload Genbank genome sequences. Users can also upload their own unpublished genome sequences. When a user attempts to upload a genome sequence assembly that is already in LINbase, users will be redirected to that genome sequence.

If the user’s genome sequence is not yet in LINbase, a LIN will be assigned using the LINflow procedure described in detail elsewhere (manuscript in preparation). In short, k-mer signatures are computed using sourmash [19], with parameters k=21 and k=51. The computed signatures are then compared with the signatures of representative genomes that are already in LINbase at the 95% ANI level (LIN position F) using k=21. If a genome sequence is found to have a Jaccard similarity of J >= 0.2475 (which corresponds to 95% ANI) compared to the uploaded genome, the uploaded genome is identified as a member of the represented LINgroup and the signature of the new genome is then compared with the signatures of all the members of this LINgroup using k=51. If instead, the LINgroup with the highest Jaccard similarity has a J<0.2475, the signature of the new genome is compared with the members of that LINgroup using k=21. In both cases, ANI is then calculated between the uploaded genome and the genome with the highest Jaccard similarity using pyANI [20]. The computed ANI value is then used to assign a LIN to the new genome based on the LIN of the genome with the highest Jaccard similarity. This is done by keeping the prefix of the reference LIN up to the LIN position at which the ANI threshold is smaller than the computed ANI value. At the next LIN position (*i*.*e*., at which the computed ANI value is smaller than the ANI threshold), a number is assigned that has not yet been used at that position. The following positions are filled with 0’s. The average time for one LIN assignment is currently 3 minutes and 54 seconds.

When uploading a genome, the user has to enter a strain name as the only required metadata value. Genus, species, and information on intraspecific classification are optional. Other metadata can be entered based on a user’s selection of “Interest” (**Table 1** and **2**). Currently, the following interests are available (but additional interests and additional metadata options can be added upon contacting LINbase administrators at): Undefined interest, Plant pathogens, Environmental bacteria, Uncultured bacteria, Foodborne pathogens, and Archaea (**Figure 3**).

**Table 1.** The “Metadata” table of LINbase. Each Metadata ID is associated with a category of metadata (Metadata_Item).

| Interest_ID | InterestName | Metadata_IDs |
| --- | --- | --- |
| 1 | Plant pathogens | 1,2,3,4,5,6,7,8,9,10,11,12,13,14 |
| 2 | Foodborne pathogens | 1,16,17,2,3,4,5,6,7,8 |
| 3 | Environmental bacteria | 1,2,3,4,5,6,7,8,15 |
| 4 | Uncultured bacteria | 1,2,3,4,5,6,7,8,15 |
| 5 | Archaea | 1,2,3,4,5,6,7,8,15 |
| 6 | Unidentified interest | 1,2,3,4,5,6,7,8,15 |

**Table 2.** The “Interest” table of LINbase. The current interests in LINbase and their corresponding metadata categories represented as lists of Metadata IDs (Metadata_IDs).

| Metadata_ID | Metadata_Item |
| --- | --- |
| 1 | Type strain |
| 2 | NCBI Taxonomy ID |
| 3 | NCBI Accession Number |
| 4 | Date of isolation |
| 5 | Country |
| 6 | Region |
| 7 | GPS Coordinates |
| 8 | Link to peer-reviewed paper |
| 9 | Host of isolation |
| 10 | Secondary host |
| 11 | Disease |
| 12 | Symptom |
| 13 | Phenotype |
| 14 | Fluorescence |
| 15 | Environmental source |
| 16 | Source of isolation |
| 17 | Outbreak |

**Figure 3.**
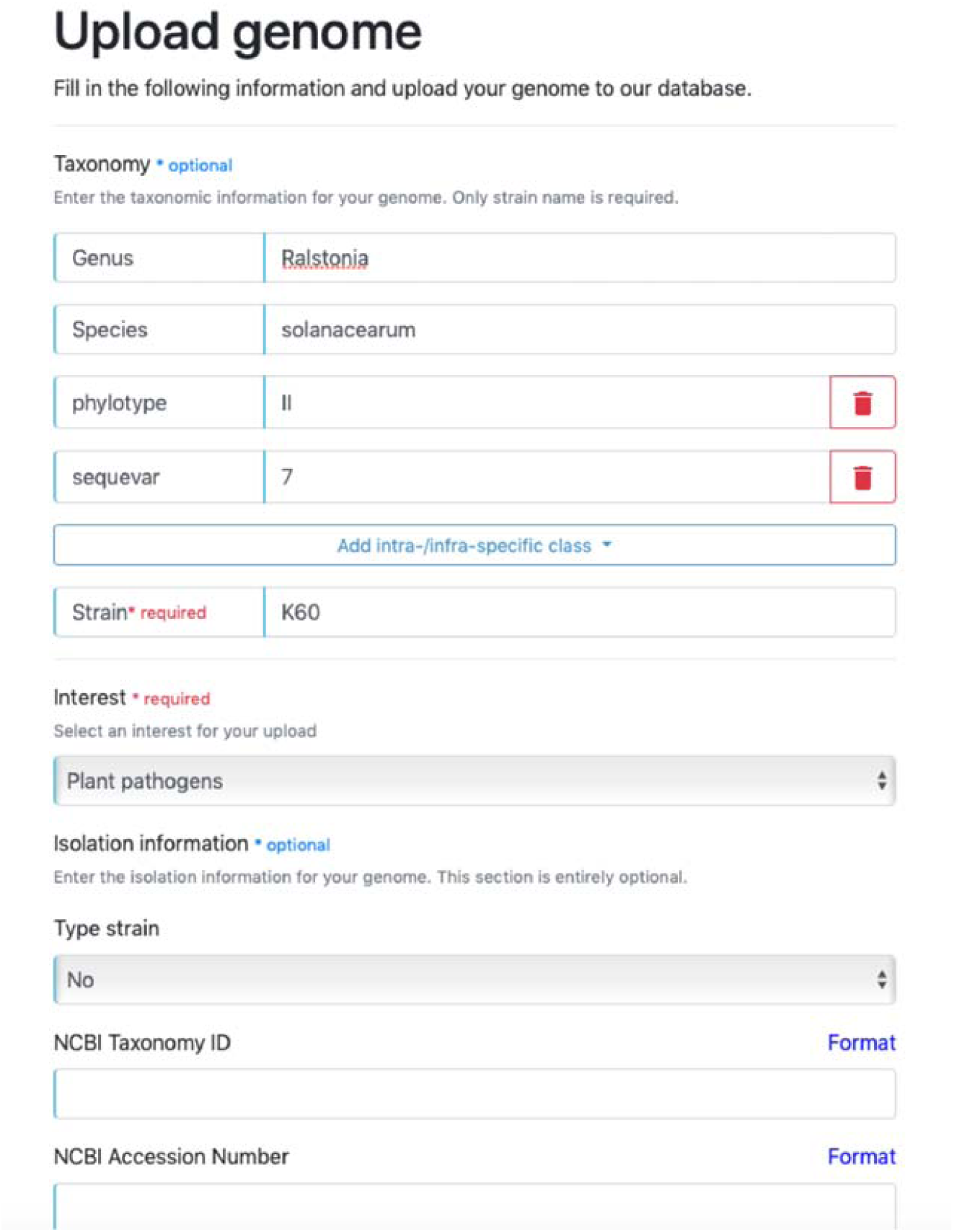
“Upload genome” form. Users are asked to enter taxonomic information and isolation metadata before uploading microbial genome sequences. In the taxonomic information section, only strain name is required as an identifier of the uploaded genome since taxonomic information may not be available. Users are required to choose an area of interest to associate the uploaded genome with a research area, e.g. Plant pathogen, Foodborne pathogen, Environmental bacteria, etc. This allows the form to change dynamically in regard to the available metadata fields. For example, the field “Host of isolation” only becomes available when choosing “Plant pathogens” but not when choosing “Environmental bacteria”.

After the genome is successfully uploaded, the result page will return the LIN assigned to the new genome, the most similar genome based on which the LIN was assigned, and the respective ANI value. The genome’s membership in LINgroup(s) that have been described in LINbase by the same user or any other user are also reported. A description of how LINgroups are described follows below.

### LINgroup description function

A group of genomes can be selected from any result page and described as a LINgroup by highlighting with the mouse the LINprefix shared by the group of genomes and clicking on the link “add a description”. The user chooses the type of LINgroup (either a taxonomic rank or a non-taxonomic group within a species that share the same characteristics), adds a name (which can be a species name, if the LINgroup corresponds to a species, or any other name the user chooses), a description giving more information about the LINgroup (for example, the phenotype that is shared by its members), and a URL or DOI to a peer-reviewed publication about the LINgroup (**Figure 4**).

**Figure 4.**
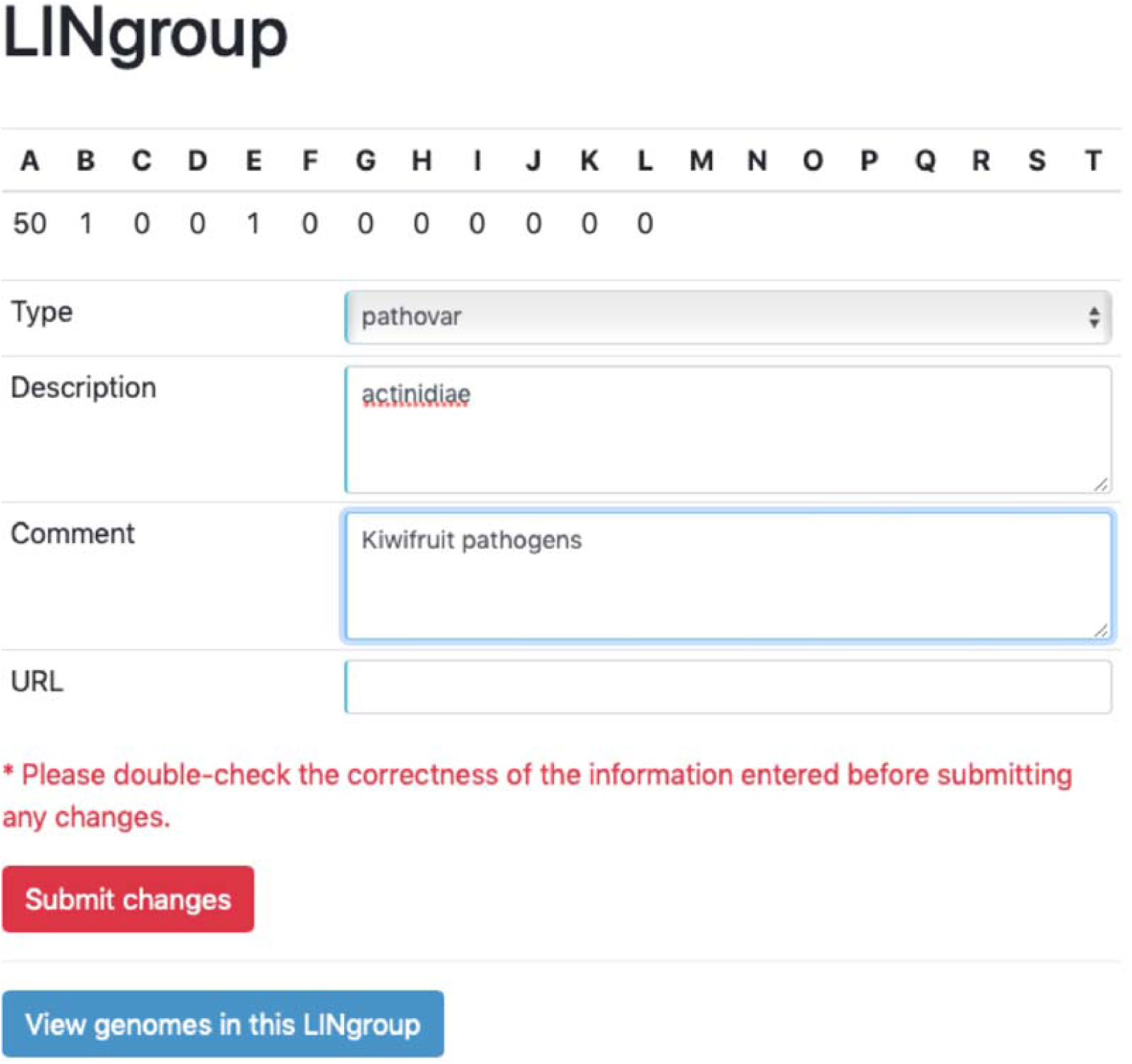
“Add a Lingroup description” form. After an undescribed LINgroup is selected, the user can describe the LINgroup at a taxonomic rank or as a group of microbes within a species that share a phenotype. This is done by choosing the type of taxon from the “Type” dropdown menu and entering a name and an optional comment and/or optional link to a peer-reviewed publication.

We expect users to generally choose the longest LIN prefix shared by a group of genomes when describing a LINgroup. For example, if all genomes of the genus *Pseudomonas* in LINbase share the LIN prefix 50_A_, then a user could describe the LINgroup 50_A_ as genus *Pseudomonas*. Instead of choosing the maximum length of the LINprefix shared by a group of genomes, a user can also choose to describe a group of genomes by the minimum length of the LINprefix that distinguishes the group from members outside of the group. For example, if there are only two genomes in LINbase that belong to an intraspecific group, this may be the better approach since more diverse members of the group may be added later. Finally, a LINgroup can also be described based on a single genome choosing the LINprefix up to position F or G, which correspond to the broadly accepted ANI thresholds for speciation at 95%-96%. This will be typically done if only the genome of the type strain of a species has been added to LINbase.

As soon as a new LINgroup description has been added to LINbase, any newly uploaded genome will be automatically identified as a member of that LINgroup if its LIN includes the LINprefix of the LINgroup.

### Genome and LINgroup search function

Both, individual genomes and described LINgroups, can be searched in LINbase. Entered parameters will form one single query so that query time is minimized. Searching by either genome or LINgroup takes less than 1 second to return the result.

When searching for genomes, users can use any LIN position(s), area of interest, taxonomic information, and isolation metadata as filters in the query (**Figure 5A**). The genome-search result page will list the genomes that match the query as well as the described LINgroups that include these genomes as members (**Figure 6**).

**Figure 5.**
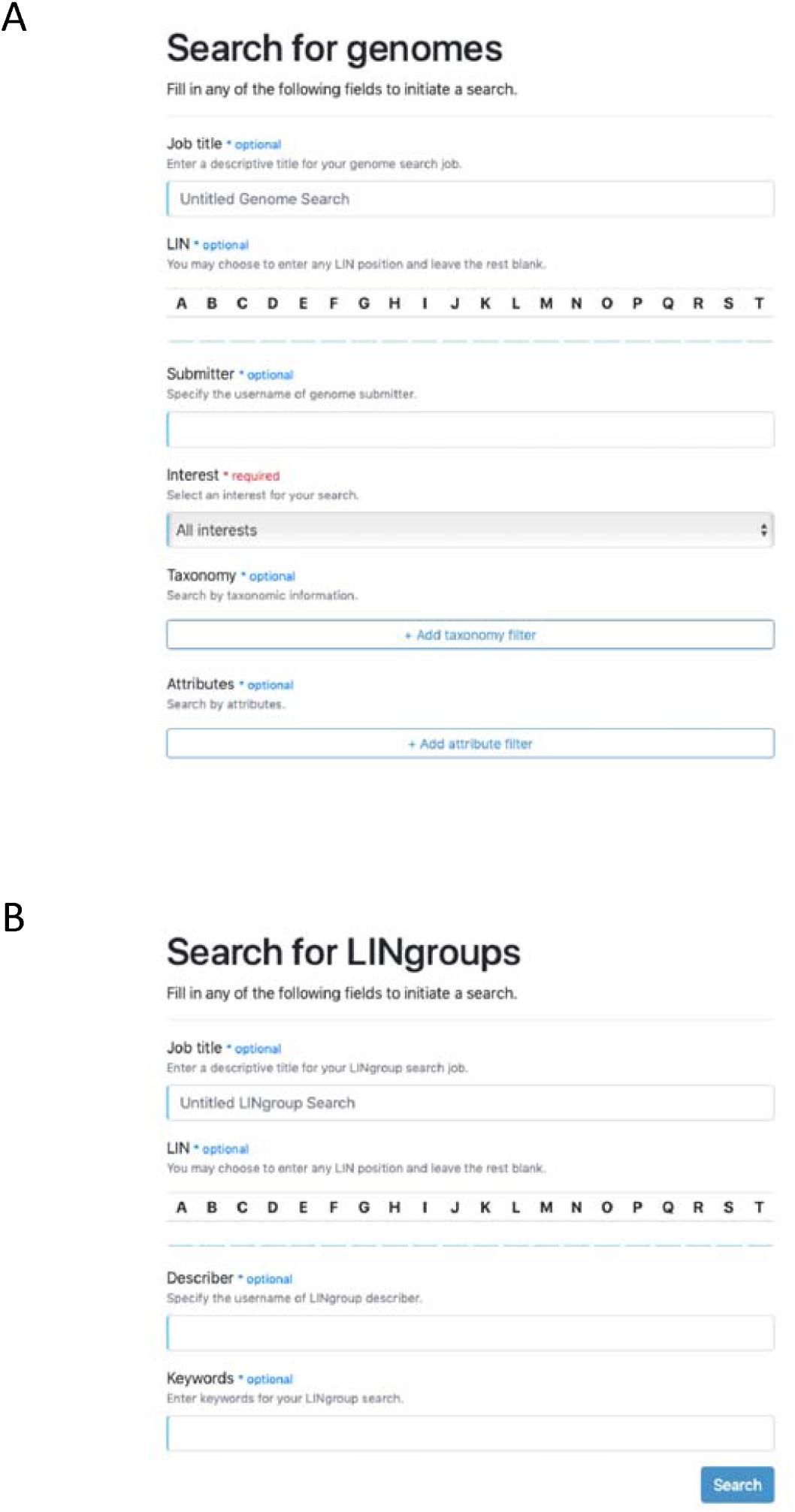
Forms for searching LINbase. (A) “Search for genomes” form. The user can search for genomes in LINbase by using any of the provided fields including LIN, submitter, interest, taxonomic information, and isolation information to narrow down the search. (B) “Search for LINgroups” form. The user can search for described LINgroups by describer and keywords. All filled fields will be passed to the backend as filters to query the database.

**Figure 6.**
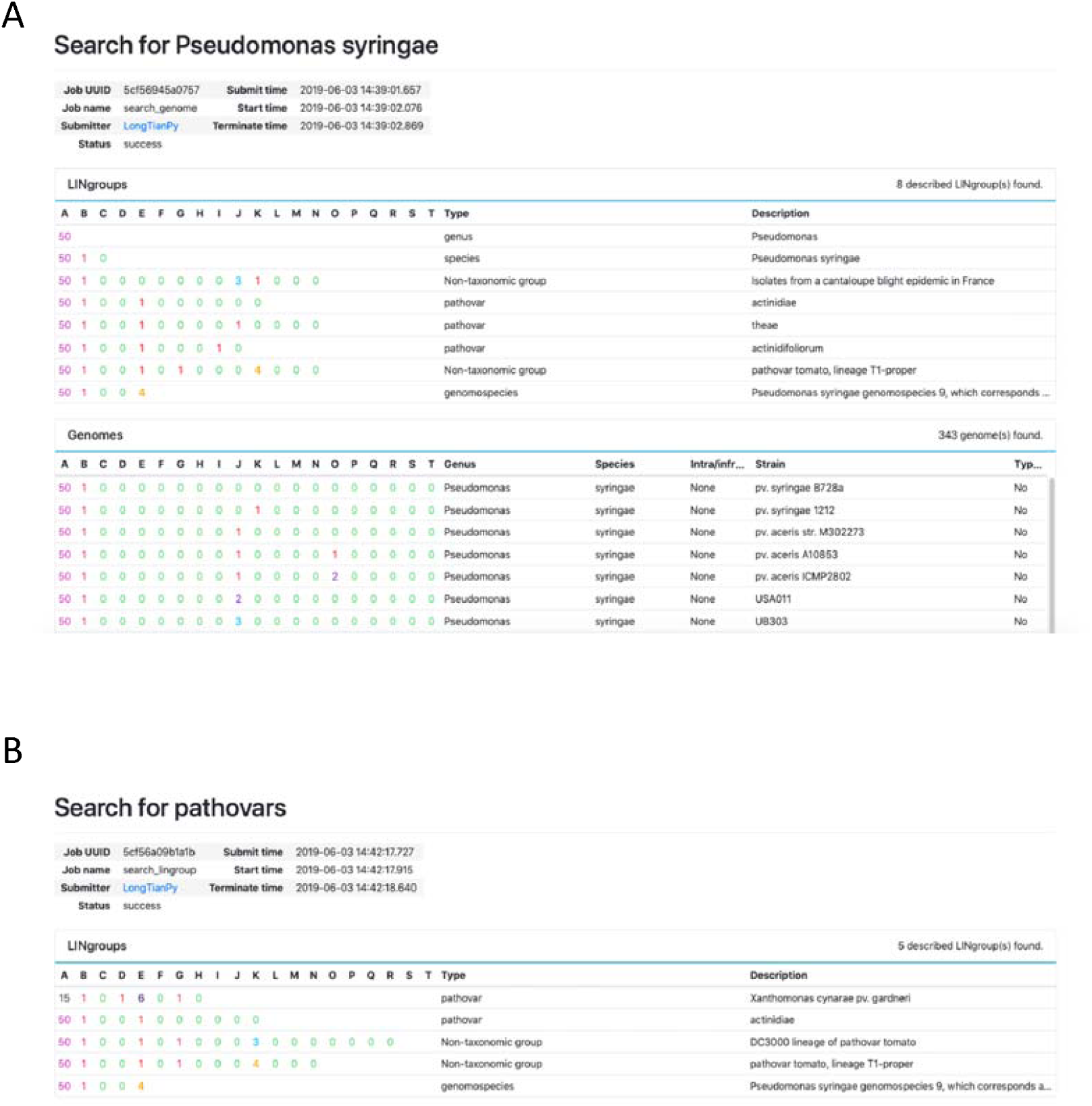
Search result pages. **(A)** The result page when searching, for example, for the species *Pseudomonas syringae*. All *Pseudomonas syringae* genomes and all associated LINgroups are printed to the screen. (B) The result page when searching, for example, for the keyword “pathovar”. All LINgroups with “pathovar” in their description are returned.

When searching for described LINgroups, users can search by LIN position(s), the name of the user who described the LINgroup, and words used in the LINgroup name and description (**Figure 5B**). The result page will list the described LINgroups that match the query.

### Identify function using gene or genome sequences as query

Users can identify an unknown microbe either using a genome sequence or a gene sequence as the query.

When using a genome sequence as the query, the most similar genome in the database is identified using a workflow similar to the one described above for the genome upload function (**Figure 7A**). However, to achieve higher speed with only moderate reduction in accuracy, FastANI [21] replaces pyANI when computing ANI between the query genome and the most similar genome identified by sourmash. On the result page, the most similar genome and its LIN, the ANI value between the query genome and the most similar genome, and any LINgroup that the query genome is a member of are reported (**Figure 8A**).

**Figure 7.**
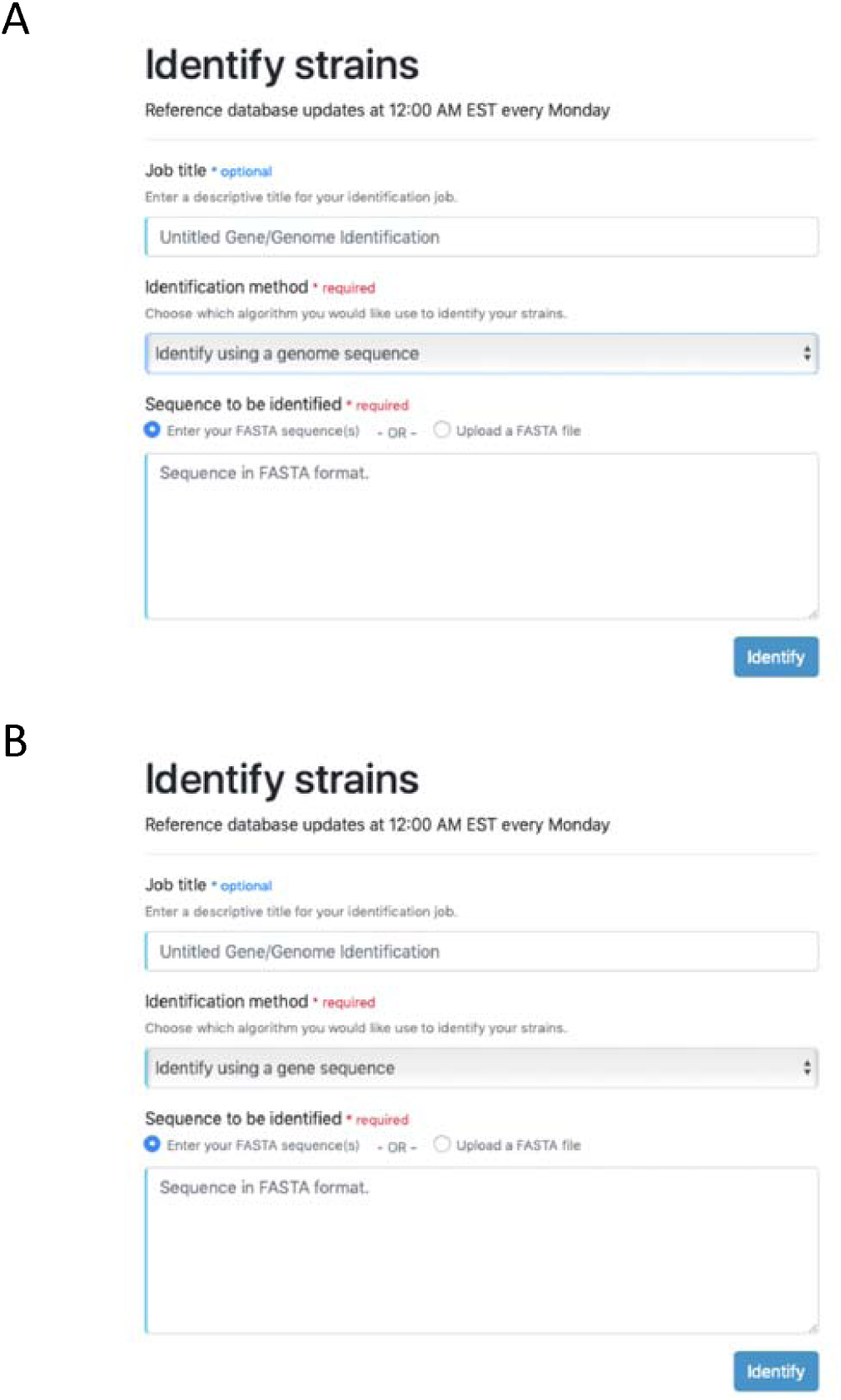
“Identify” forms. (A) Identification with a genome assembly. (B) Identification with a gene sequence. Both functions accept gene or genome sequences uploaded as a FASTA-format file or entered in the textbox.

**Figure 8.**
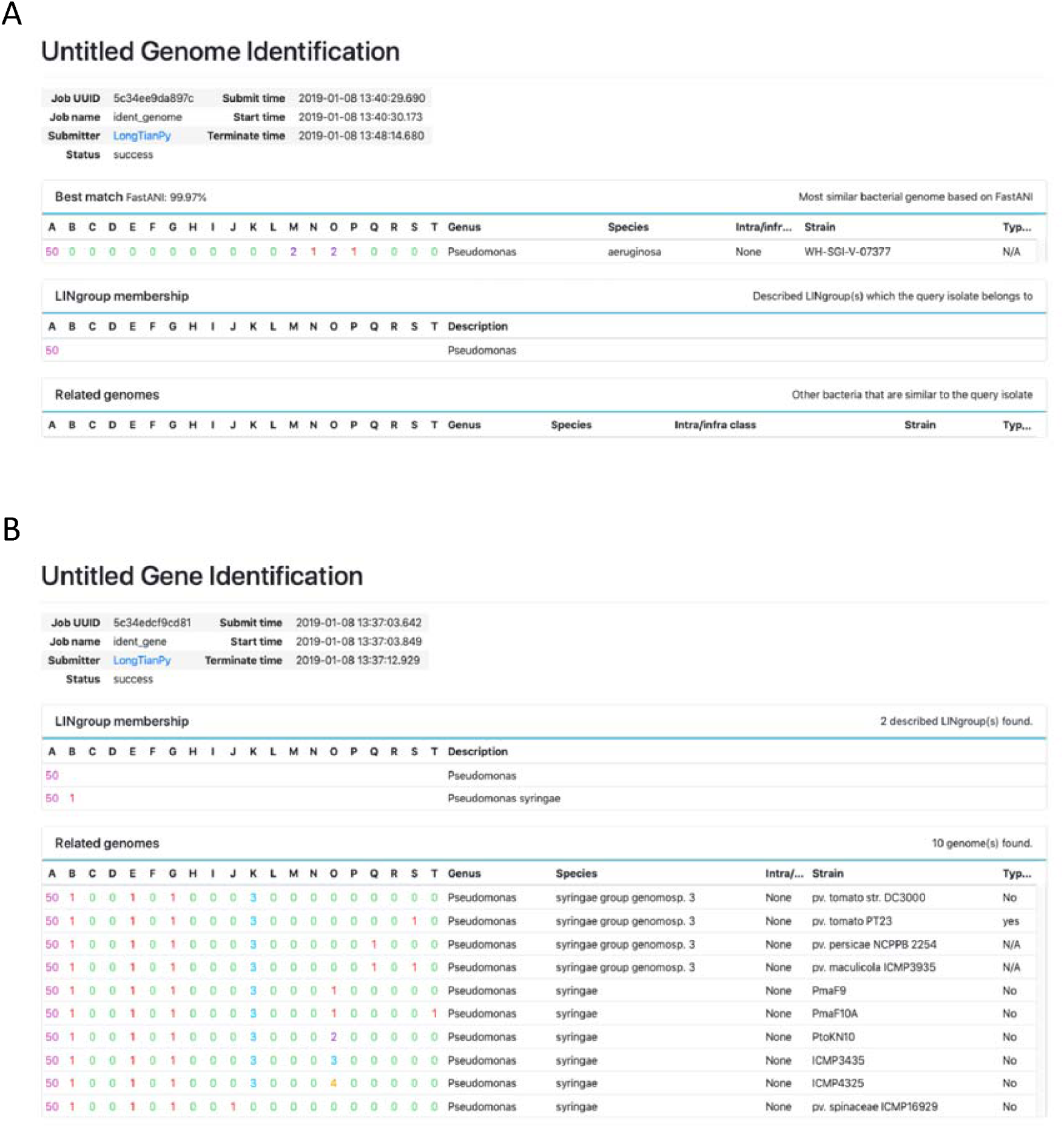
“Identify” result pages. (A) Result page for genome-based identification. The submitted genome is queried against LINbase genomes and the genome with the highest FastANI is returned. The LINgroups that the query genome belongs to that are based on its ANI with the best match are listed as well. (B) Result page for gene-based identification. The submitted gene sequence is queried against LINbase genomes with BLASTN. Genomes with E-value = 0 are listed as best matches. The LINgroups the best matches belong to are listed as well. For both types of identification, the submitted sequences will be deleted from the server once the query is completed.

Users who do not have the whole genome sequence of an unknown microbe can also use a single gene sequence as the query in combination with BLASTn (**Figure 7B**) [22]. However, accuracy is of course largely reduced, since multiple genomes, which may even belong to different LINgroups, may align with a short gene sequence with 100% identity. To minimize the risk of misidentification, only genomes with low e-values are returned on the result page along with LINgroup(s) that these genomes belong to (Figure 8B).

### Comment function

A commenting system is implemented in the LINgroup profile page to facilitate communication and potential collaboration among LINbase users. Users can add comments to any LINgroup (described or undescribed) to discuss the LINgroup with other users. Posted comments can be edited or deleted by the original poster. At this time, users are not automatically notified of comments posted to LINgroups they described. However, this function is planned for the future.

## Data security and dissemination

Genome assemblies in LINbase, either sourced from public databases, such as NCBI, or uploaded by users, are securely saved on the server and cannot be viewed or downloaded by any user. Gene and genome sequences uploaded as part of the identification function are deleted along with intermediate data immediately after the identification process is finished. The data that are shared in LINbase are genome metadata (including taxonomic and isolation information), LINs, LINgroups, LINgroup descriptions, and comments. Therefore, LINbase is ideally suited for sharing the precise identity of sequenced genomes as soon as they are generated while keeping the actual genome sequences private until submission to a public database.

## Discussion

Here we introduced LINbase, a Web service that implements bacterial taxonomy based on whole genome similarity and supported by fast and accurate algorithms. LINbase complements functionalities offered by other online Web services for genome-based microbial identification, such as MiGA [23] or EzBioCloud [24], as follows: 1. it labels individual genomes with LINs, which reflect the precise genomic relatedness among strains in the database; 2. it automatically gathers genomically similar bacteria into taxa (LINgroups); 3. it provides a user-friendly interface to genomically circumscribe validly published named taxa at the genus and species rank and at intraspecific levels as LINgroups permitting precise genome-based identification; 4. it uses crowdsourcing to incorporate informal taxa/LINgroups independently of published named taxa; 5. it encourages scientific exchange and early sharing of data by providing an avenue to share the precise identity of sequenced genomes without sharing the genome sequences themselves; and 6. it allows users to interact with each other by commenting on LINgroup circumscriptions and descriptions.

Despite the aforementioned advantages of LINbase, there are limitations in its current version in regard to the classification of bacteria at higher ranks (family, order, class, and phylum), which can currently not be circumscribed as LINgroups, and for bacteria with very recent common ancestors, *e*.*g*., differentiating foodborne pathogens from different outbreaks is currently only possible when high quality genome assemblies are available. If assemblies are of low quality, the correlation between phylogeny and LINs fails at the right-most LIN positions. Also, genome upload is currently managed by a scheduler that only allows one process at a time. This limits the ability to batch upload genomes and does not allow multiple users to upload genomes at the same time.

Future implementations of LINbase will focus on increasing the speed of the identification function when using a genome sequence as the query and of LIN assignment. Parallelization is a promising solution to speed up LIN assignment when genomes are uploaded by different users at the same time. Parallelization would also allow batch uploading, which can further accelerate identification and LIN assignment. We are also planning to expand LIN positions to the left up to the phylum level by using algorithms to detect low-level genome similarity and to improve assignment of isolates to outbreaks by integrating additional algorithms to precisely identify phylogenetic relationships among very similar genomes. Finally, our aim is to automatically add all genome sequences in Genbank to LINbase, to automatically circumscribe all monophyletic taxa as LINgroups by integrating LINbase with the Genome Taxonomy Database [25], and to integrate LINbase with other platforms to improve genome-based classification and identification of microbes at all taxonomic ranks.

## Author contributions

BAV and LSH conceived the initial idea of LINbase. LT is the main author of LINbase. CH took part in the development of the front end of LINbase and the optimization of the database. LT, LSH and BAV prepared the manuscript. All authors read and approved the final version of the manuscript.

## Acknowledgements

The authors would like to acknowledge the undergraduate students of Computer Science Department at Virginia Polytechnic Institute and State University, Grant Hughes, Vincent Eastman and Teresa Paul for participating in the early design of the user interface.

